# Unconventional secretion of angiogenic sonic-hedgehog-containing large extracellular vesicles is stimulated via the PI3K–Rab18-GDP pathway

**DOI:** 10.1101/2025.09.06.674659

**Authors:** Shuo Wang, Rio Imai, Yosuke Tanaka

## Abstract

Large extracellular vesicles (lEVs), with diameters >600 nm, play essential roles in special types of intercellular communications. However, information on the molecular mechanism underlying their roles and on their relevance remains scarce. Here we show that PI3K–Rab18-GDP signaling facilitates the secretion of lEVs from human mesenchymal stem cells (hMSCs) and fibroblasts. These EVs contain a large amount of sonic hedgehog (SHH) protein and promote angiogenesis. Rab18 serves as the master facilitator of this secretion only in its GDP-bound form, which can be enriched by the Rab inhibitor CID1067700 or by the PI3K agonist SF1670. Rab18-GDP accumulates in the cell center and develops SHH-lEV precursors from perinuclear endosomes. Rab18-GDP recruits heat shock protein 90α (Hsp90α) and neutral sphingomyelinase 2 (nSMase2) and facilitates vertical secretion of SHH-lEVs with an Hsp90α corona from the cell center. This secretory machinery may play an important role in developmental and regenerative PI3K-mediated angiogenesis.

## 1. Introduction

Extracellular vesicles (EVs) carry proteins, lipids, and nucleotides as freight between tissues and alter the functional properties of recipient cells (Abels and Breakefield, 2016; Buzas, 2023; Veziroglu and Mias, 2020). This intercellular transfer facilitates cell–cell communication in multicellular organisms and shapes the morphogen gradients in developing tissues (Teleman et al., 2001; Wu et al., 2022; Wang et al., 2022; Tanaka et al., 2023). EVs derived from human mesenchymal stem cells (hMSCs) have a high potential for use in regenerative medicine (Musiał-Wysocka et al., 2019; Nakazaki et al., 2021), and can minimize the risk of infection and immune rejection associated with conventional allogeneic MSC transplantations (Nakazaki et al., 2024; Zhang et al., 2020). Furthermore, loading of the morphogen, sonic hedgehog (SHH), to hMSCs can increase the clinical potency of the hMSC-EVs in ischemic heart failure (Mackie et al., 2012). However, mechanisms underlying the formation of this clinically relevant EV subclass have not been determined in detail.

We previously characterized a PI3K-induced pathway for the biogenesis of SHH-containing large EVs (SHH-lEVs) based on molecular and cellular analyses of *Kif3b^LacZ/LacZ^* polydactyly mouse embryos (Wang et al., 2022). According to a distal-to-proximal FGF/PI3K signaling gradient of the limb bud mesenchyme, the secretion of SHH-lEVs and their transcytosis is locally enhanced in the peripheral layer. This mutually exclusive difference in the cellular behavior between the two layers may underlie the formation of a sustainable SHH gradient along the peripheral layer that is essential for proper digit patterning. We could reproduce this PI3K-dependent secretion of SHH-lEVs in SHHN-EGFP-transfected fibroblasts. As evident from fluorescent microscopy, exceptionally large fluorescent particles were secreted from the cells, which may be classified as a species of lEVs. SHH-lEV precursor organelles were associated with “ART-EV” markers, Axl, Rab18, and Tmed10 (Coulter et al., 2018; Wang et al., 2022). However, the clinical potency and mechanism of SHH-lEV biogenesis have remained elusive.

Rab18 belongs to the Rab GTPase superfamily, the members of which are molecular determinants of membrane organelle trafficking within the cytoplasm (Schäfer et al., 2000). The conformation of Rab proteins alters significantly between the GTP- and GDP-bound states, which are associated with specific effector proteins (Homma et al., 2021; Klopper et al., 2012). In particular, distinct preferences of the GTP- and GDP-bound forms of Rab proteins for plus- and minus-end-directed molecular motors result in the dispersal or accumulation of Rab-bound organelles, respectively (Niwa et al., 2008; Ueno et al., 2011; Wanschers et al., 2008). Rab18 is a binding partner of kinectin, which is a protein associated with kinesin-1 light chain (KLC) in the ER (Guadagno et al., 2020). Because kinesin-1 is the initial member of the plus-end-directed kinesin superfamily of molecular motors that significantly regulates the organelle distribution and ER function (Tanaka et al., 2024; Tanaka et al., 1998), the dynamic interaction– dissociation kinetics between Rab18 and kinesin-1 may play an essential regulatory role in SHH-lEV biogenesis.

Hsp90α is a multifunctional heat shock protein. It generates cytoplasmic condensations with kinesin-1 and is involved in chaperone exchange in the ER sheets that helps proper folding of calcium channels and prevents diabetes (Tanaka et al., 2024). As its distinct function, Hsp90α facilitates membrane deformation for exosome biogenesis (Lauwers et al., 2018). It is also extracellularly secreted and believed to stabilize extracellular enzymes such as metalloproteinases and to facilitate metastatic niche formation (Baker-Williams et al., 2019).

In this study, we sought to test if the Rab inhibitor CID1067700 (CID) affects the secretion of small EVs (sEVs) or lEVs. We identified that it significantly and specifically enhanced SHH-lEV secretion. We provide evidence for a strong angiogenic potency of SHH-lEVs, and describe the molecular mechanism of PI3K–Rab18-GDP-mediated SHH-lEV biogenesis in detail. PI3K signaling enriches the GDP-bound form of Rab18, which accumulates in perinuclear endosomes because of kinesin-1 dissociation. They develop SHH-lEV precursors via recruiting and activating the Rab18-GDP effector proteins, including Hsp90α and neutral sphingomyelinase 2 (nSMase2). After 1 h of CID stimulation, they are secreted exclusively from the perinuclear region. After 24 h, they further form extracellularly floating Hsp90α condensations containing SHH-lEVs on cytonemes. These findings regarding Rab18-GDP-mediated SHH-lEV secretion will facilitate translational research on SHH-lEVs that should be useful in regenerative medicine, and open the door for understanding the cell biology of unconventional secretion of lEVs.

## 2. Materials and Methods

### 2.1 Cells

hMSCs from bone marrow were purchased from Lonza (Cat. No. PT-2501), and they were cultured and characterized according to the manufacturer’s protocols. Briefly, they were routinely cultured in MSCBM™ Mesenchymal Stem Cell Basal Medium (PT3238, Lonza), supplemented with MSCGM SingleQuots™ Supplements and Growth Factors (PT4105, Lonza), in 10 cm dishes (#150466, Thermo Scientific). They cells were detached by treating with trypsin/EDTA solution (CC-3232, Lonza) for 5 min at 37°C, seeded at a density of 5,000–6,000 cells/cm^2^ every 7 days for successive passages, and used within six passages. The culture medium was changed every three days.

Human umbilical vein endothelial cells (HUVECs) were purchased from Lonza (C-2519AS) and routinely cultured in EBM™-2 Endothelial Cell Growth Basal Medium-2 (CC-3156, Lonza) with EGM™-2 SingleQuots™ Supplements (CC-4176, Lonza), following the manufacturer’s protocols. For passaging, after rinsing with HEPES buffer (CC-5022, Lonza), HUVECs were detached by treating with trypsin/EDTA (CC-5012, Lonza) for 3–5 min at 37°C, harvested using a trypsin neutralizing solution (CC-5002, Lonza), and seeded at a density of 2,500 cells/cm^2^ every 7 days for successive passages. They were used within four passages, and the culture medium was changed every three days. At each passage, HUVECs were plated into an eight-chambered LabTek II Chambered #1.5 German Coverglass System (#155409, Thermo Scientific) precoated with Matrigel (#356231, Corning) and used for angiogenesis assays of the hMSC-EVs. NIH3T3 cells were routinely cultured in high glucose DMEM containing 10% FBS (F7524, JRH Biosciences), according to standard procedures. The cells were stably transfected with EGFP-Rab18 plasmids or Piggybac-SHHN-mStayGold expression vector with a hyPBase expression vector (VB010000-9365tax, Vector Builder) using Invitrogen Lipofectamine LTX with PLUS Reagent (#15338100, Thermo Fisher), containing 200 μg/mL G418 (#10131035, Thermo Fisher) or 15 μg/mL puromycin (P8833, Sigma Aldrich), respectively, and subcloned for cell biological assays.

### 2.2 Pharmacology

CID1067700 was purchased from Sigma-Aldrich (SML0545), solubilized at 40 mM in DMSO, and applied to cells at 40 µM overnight. TAS116 was purchased from MedChemExpress (HY-15785), solubilized at 1 mM in DMSO, and applied to cells at 0.5 µM for 48 h. GW4869 was purchased from Cayman Chemical (#13127), solubilized at 5 mM in DMSO, and applied to cells at 1.25 µM overnight. SF1670 (S7310, Selleck) was used at 250 nM, as previously described (Tanaka et al., 2016).

### 2.3 EV secretion

The handling of EVs was performed according to the MISEV2023 criteria (Welsh et al., 2024). hMSCs were cultured in 6-well plates or 10-centimeter dishes at a density of 5,000–6,000 cells/cm^2^ for 6 days and stimulated with SF1670 (250 nM), CID1067700 (40 µM), or DMSO (1/1,000) in the EV-Up medium (#053-09451, FUJIFILM-Wako) for 1–24 h. The conditioned medium was collected, centrifuged at 200–500 *× g* for 10– 30 min at 4°C to remove the cell debris, optionally condensed to 1 mL with a Millipore Amicon Ultra 30 kDa MWCO filter (MAP030C37, Pall), and subjected to analyses. For concentrating secretomes, the conditioned medium was mixed with 1/2 volume of Total Exosome Isolation Reagent (Thermo Fisher), incubated overnight at 4°C with rotating, and centrifuged at 10,000 *× g* for 60 min at 4°C using a bench-top centrifuge (TOMY Seiko). The pellets were resuspended in PBS or PBS-HAT medium (Görgens et al., 2022), stored at 4°C, and used in assays within 2 weeks.

### 2.4 Antibodies

A rabbit anti-SHH antibody (20697-1-AP, RRID: AB_10694828, 1:1,000) was purchased from Proteintech. A rat monoclonal anti-Hsp90 antibody 16F1 (ADI-SPA-835, RRID: AB_311881, 1:500) was purchased from Enzo Life Sciences. A mouse monoclonal anti-TSG101 antibody 4A10 (MA1-23296, RRID: AB_2208088, 1:1,000) was purchased from Thermo Fisher. A rabbit monoclonal anti-CD81 antibody D5O2Q (Cat. No. 10037, RRID: AB_2714207) was purchased from Cell Signaling Technology. A rabbit anti-glutathione S-transferase (GST) antibody (A5838, RRID: AB_258261, 1:1,000), mouse anti-gamma-adaptin (AP-1) monoclonal antibody (A4200, RRID: AB_476720, 1:1,000), mouse anti-assembly protein 180 (AP-3) monoclonal antibody (A4825, RRID: AB_258203, 1:1,000), and mouse IgM anti-ceramide antibody (C8104, RRID: AB_259087, 1:500) were purchased from Sigma-Aldrich. A rabbit anti-KIF5A N-terminal antibody (RRID: AB_2571744, 1:1,000) was described previously (Kanai et al., 2000).

Among the secondary antibodies, horseradish-peroxidase (HRP)-conjugated anti-rabbit and mouse IgG antibodies (1:1,000) were purchased from GE Healthcare Life Sciences; alkaline-phosphatase-conjugated anti-rabbit and mouse IgG antibodies (1:1,000) were from Cappel; Alexa-Fluor 405-, 488-, and 568-conjugated anti-mouse, anti-rat, and anti-rabbit IgG and anti-mouse IgM antibodies (1:200–1:500) were from Thermo Fisher.

### 2.5 Expression Vectors

SHHN-EGFP expression vectors (plasmid, adenovirus) for amino acids 1–198 of the N-terminus of mouse SHH were previously described (Wang et al., 2022). For constructing the SHHN-tagRFP expression vector, the *Shh-N* cDNA was amplified via PCR using the primers *Sal-SHHATGKozak-fwd* (5′-ACCGTCGACCATGGTGCTGCTGCTGGCCAGATG-3′) and *Bam-SHH198-rev* (5′-CAAGGATCCCGGCCGCCGGATTTGGCCGCCACG-3′), and ligated with the *pTagRFP-N* vector (FP142, Evrogen) to generate the *pSHHN-tagRFP* expression plasmid.

To construct the Hsp90-tagRFP expression vector, mouse *Hsp90aa1* cDNA was amplified from FANTOM clone I1C0020P08 (Kawai et al., 2001) via PCR using the primers Xho+hsp90aa1atgFw (5′-ACCCTCGAGCTATGCCTGAGGAAACCCAGACCCAAG-3′) and Kpn+hsp90 aa1stopRev (5′-ACCGGTACCTTAGTCTACTTCTTCCATGCGT-GATGTGTC-3′), and ligated into the *pTagRFP-C1* vector at XhoI and KpnI sites. The inserted cDNA was ligated into the *pEGFP-C1* (Clontech) vector. These expression units were transferred to the ViraPower Adenoviral Expression System (Thermo Fisher) according to the manufacturer’s instructions.

EGFP-Rab18Q67L (Addgene plasmid #49596; https://www.addgene.org/49596; RRID: Addgene_49596), EGFP-Rab18S22N (Addgene plasmid #49597; https://www.addgene.org/49597; RRID: Addgene_49597), and EGFP-Rab18 (Addgene plasmid #49550; https://www.addgene.org/49550; RRID: Addgene_49550) expression vectors were gifts from Marci Scidmore (Huang et al., 2010). For expression of GST-tagged Rab18 proteins in *Escherichia coli*, the Rab18 fragment of each plasmid was amplified via PCR using the following primers: EcoRI+Rab18-F (5′-TCCGAATTCCATGGACGAGGACGTGCTGACCACTCTG-3′) and Not+Rab18-R (5′-AATGCGGCCGCCTATAGCACAGAGCAGTAACCGCCGCAGGCG-3′) and ligated with the *pGEX-4T-3* vector (Cytiva Life Sciences) at EcoRI and NotI sites.

The piggybac-SHHN-mStayGold expression vector was constructed by Yiming Li. Briefly, the *Shh-N* cDNA was amplified via PCR from the *pSHHN-EGFP* vector (Wang et al., 2023) using the primers newPiggymShhN-F (5′-TCAGATCCGCTAGCGCCACCATGGTGCTGCTGCTGGCCAG-3′) and newPiggymShhN-R (5′-ACCATTGATCCGCCGCCACCGCCGCCGGATTTGGCCGC-3′). The vector backbone was amplified from a PiggyBac-based expression vector, with a *pPB[Exp]-Puro* backbone (Vector Builder, VB240319-1202jdr), using primers piggybac-F (5′-ACCCAGCTTTCTTGTACAAAGTG-3′) and piggybac-R (5′-CAACTTTTCTATACAAAGTTGATGATAT-3′). The CMV promoter was amplified from *pEGFP-N1* (Clontech) using primers CMV-F (5′-AACTTTGTATAGAAAAGTTGCGTTACATAACTTACGGTAAATGG-3′) and CMV-R (5′-GCTAGCGGATCTGACGGTT-3′). The fluorescent protein *mStayGold* cDNA was amplified from its ER expression vector (kindly provided by Dr. Atsushi Miyawaki, RIKEN) (Hirano et al., 2022) using primers SG-F (5′-GGTGGCGGCGGATCAATGGTGAGCAAGGGCGAGGA-3′) and SG-R (5′-TTTGTACAAGAAAGCTGGGTTTACAGGTGGGCCTCCAGGGT-3′).

All four PCR products were purified using the QIAquick Gel Extraction Kit (QIAGEN, Cat. No. 28704) and assembled using the Gibson Assembly Master Mix (E2611, New England Biolabs) at 50°C for 60 min. The entire ligation mixture was added to 50 µL of *E. coli* DH5α competent cells, incubated on ice for 30 min, subjected to heat shock at 42°C for 45 s, and then incubated on ice for 2 min. After adding 950 µL of antibiotic-free LB medium, the cells were shaken at 250 rpm for 1 h at 37°C. Thereafter, 500 µL of cell suspension was plated on LB agar containing ampicillin and incubated overnight at 37°C. Next day, five colonies were picked and cultured in 5 mL LB medium with ampicillin. After 2 h, colony PCR was performed to verify plasmid incorporation. Two positive clones were further cultured for 12 h, and plasmids were extracted using the QIAprep Spin Miniprep Kit (QIAGEN, Cat. No. 27104). Plasmids were sent to Eurofins Genomics for Sanger sequencing. The hyPBase expression vector *pRP[Exp]-mCherry-CAG>hyPBase* was purchased from Vector Builder (Cat No. pDNA(VB010000-9365tax)-P).

### 2.6 Transfection and Transduction

The expression vectors were introduced into cells either by lipofection of plasmids using Lipofectamine LTX-PLUS reagent (Thermo Fisher) or via transduction of adenoviral vectors using the ViraPower system (Thermo Fisher), according to the manufacturers’ protocols.

### 2.7 Nanoparticle Tracking Analysis (NTA)

The conditioned medium was cleared via centrifugation, diluted 1:50 with PBS, and subjected to a Zetaview Nanoparticle Tracking Analysis Instrument, selecting the “highest” mode (Particle Metrix). The collected data were analyzed and illustrated using the Prism software (v.7 and v.9.3.1, GraphPad).

### 2.8 FACS Analysis

NIH3T3 cells were stably transduced with the *piggybac-SHHN-mStayGold* expression vector and stimulated with 40 μM CID for 1 h. The conditioned medium was cleared via centrifugation, directly subjected to Beckman Coulter CytoFLEX flow cytometer (Cat. No. B53018), and the fluorescence intensity of each particle (excitation: 488 nm; emission: 525 nm) against the particle size was plotted. For estimating the particle size, 300-, 600-, and 1,000-nanometer polystyrene beads were used. Because of the difference in refractive index, the values on the *y-*axis estimated from the beads were assumed to be a little smaller than the actual values for the EVs. The region of highly fluorescent large particles (P1, **Fig. 3F–H**) was gated and compared between the transfected and non-transfected cells.

### 2.9 Transmission Electron Microscopy

Transmission electron microscopy (TEM) was performed following standard procedures, as described previously (Nonaka et al., 1998). The total EV pellets were fixed in the half-Kalnovsky fixative [2% paraformaldehyde, 2.5% glutaraldehyde, 0.1M cacodylate buffer pH 7.4] at 37°C for 10 min and then at 4°C overnight, followed by post-fixation with 1% OsO_4_ in 0.1 M cacodylate buffer (pH 7.4) on ice for 1.5 h. They were then stained with 1% uranyl acetate for 1 h and dehydrated by passing through a graded ethanol series. Finally, the pellets were embedded in Quetol-812 (Nisshin EM), cut into ultrathin sections, electron stained with uranyl acetate and lead citrate, and observed using a JEM2000 microscope at 100 keV and a JEM-1400 Flash microscope at 80 keV (JEOL).

### 2.10 Angiogenesis Assay

Angiogenesis assays were performed according to a protocol provided by Lonza Corporation (Arnaoutova and Kleinman, 2010). Briefly, eight-well glass-bottomed chambers (#155409, Thermo Scientific) were precoated with cold Matrigel (#356231, Corning) at 250 µL/well and incubated at 20°C for 10 min and at 37°C for at least 30 min in a 5% CO_2_ atmosphere. HUVECs were seeded at a density of 6.5–8 × 10^4^ cells/cm^2^ in each well of the Matrigel-coated chambers and mixed with the isolated total EV fractions from hMSCs. After 90 min, they were imaged using an LSM780 confocal microscope (ZEISS) equipped with a Plan-Apochromat 10×/0.45 M27 lens and 488 nm laser in the differential interference contrast (DIC) mode at 37°C in a 5% CO_2_ atmosphere. The data collected were analyzed using the ImageJ software (v. 1.53q) (Schindelin et al., 2015) and the number of tubes, number of branching points, and total tube length were quantified (Fujio et al., 2017; Lin et al., 2019).

### 2.11 Immunoblotting

Immunoblotting was performed as described previously (Tanaka et al., 2016). For sample preparation, EVs, purified as described above, were lysed with a 1/3 volume of 4 × Laemmli sample buffer and boiled at 98°C for 1–2 min. They were then subjected to SDS-PAGE, transferred onto a PVDF membrane (Immobilon-P, EMD Millipore), incubated with primary and HRP-conjugated secondary antibodies using Can Get Signal Solutions 1 and 2 (TOYOBO), and subjected to an electrochemiluminescence (ECL) procedure (GE Healthcare Life Sciences). Immunoreactive bands were detected using an ImageQuant LAS 4000 mini system (GE Healthcare Life Sciences) and quantified using the ImageJ software. Alternatively, they were probed with alkaline-phosphatase-conjugated secondary antibody (Cappel), and color was developed using BCIP and NBT (Roche), as described previously (Kondo et al., 1994). Molecular weight markers were purchased from Bio-Rad.

### 2.12 GST pulldown assay

GST pulldown assays were performed as described previously (Niwa et al., 2008), using B-PER Bacterial Cell Lysis Reagent (Thermo Fisher Scientific) for bacterial harvesting.

### 2.13 Density gradient centrifugation

Optiprep (iodixanol) density differential centrifugation of EVs was performed as described previously (Kowal et al., 2016). The conditioned medium was cleared by centrifuging at 800 *× g* for 10 min, and the EVs were pelleted via another round of centrifugation at 10,000 *× g* for 60 min. They were resuspended in 3 mL of 30% Optiprep (Serumwerk Bernburg AG), 250 mM sucrose, 10 mM Tris [pH 8.0], and 1 mM EDTA [pH 7.4]. Three milliliter of the sample, 0.9 mL of 20%, 0.8 mL of 10%, or 0.8 mL of 0% Optiprep in 250 mM sucrose, 10 mM Tris [pH 8.0], and 1 mM EDTA were sequentially added. The solution was centrifuged at 350,000 *× g* for 60 min at 4°C in a Sw55Ti rotor using an Optima XL-100K ultracentrifuge (Beckman Coulter) and 500 μL fractions were collected. They were diluted with 1 mL PBS, pelleted by centrifugation at 100,000 *× g* for 30 min at 4°C, and subjected to immunoblotting.

### 2.14 Fluorescence Microscopy

For light microscopy, cells were plated in 35-millimeter glass-bottomed dishes (D11130H, Matsunami), precoated with poly-L-lysine for 1 h (P4707, Sigma, 1:1,000). Immunofluorescence labeling was performed using the antibodies described above. Briefly, cells were fixed with 2% paraformaldehyde/0.1% glutaraldehyde/PBS at 37°C for 10 min, permeabilized with 0.1% Triton X-100/PBS at 20°C for 5 min, incubated overnight with primary antibody at 4°C and with the secondary antibodies for 1 h at 20°C followed by three 5-minute PBS washes, and then subjected to a spinning disk confocal microscope (Yokogawa/ZEISS) or an LSM780 laser-scanning confocal microscope with Airyscan (ZEISS), as mentioned above. Three-dimensional (3D) reconstructions were generated using the Imaris 8 software (Bitplane). The line profile was measured using the ImageJ software.

### 2.15 *In Vitro* SHH-lEV Imaging

To monitor the density of extracellular SHH-lEVs (**Fig. 3A–E**), NIH3T3 cells were plated at 1 × 10^4^ cells/well in an eight-chambered LabTek II Chambered #1.5 German Coverglass System (Thermo Fisher), precoated with poly-L-lysine (Sigma Aldrich). The cells were transduced with an adenoviral SHH-N-EGFP expression vector at days in vitro (DIV) 3. The medium was replaced with a complete medium, with or without chemicals at DIV4. After 24 h, time-lapse imaging using a spinning disk microscope (Yokogawa/ZEISS) (Tanaka et al., 2016) was conducted at 5 s intervals in 5% CO_2_ at 37°C. Finally, green fluorescent particles showing Brownian movement were identified using the Imaris 8 software (Bitplane) and quantified.

### 2.16 Live Imaging

For live imaging, cells were transduced with the fluorescent-protein-tagged vectors. EGFP-tagged Rab18 WT/SN/QL vectors were introduced into the cells using Lipofectamine LTX (Thermo Fisher) and selected on 200 μg/mL G418 (Thermo Fisher). SHHN-tagRFP was introduced into the cells using Lipofectamine LTX. Hsp90α-tagRFP and SHHN-EGFP were introduced via adenovirus. They were imaged using a spinning disk microscope CSU-W1 (ZEISS-YOKOGAWA), confocal laser-scanning microscope LSM780-Airyscan (ZEISS), or ZEISS Lattice Lightsheet 7 microscope with time-lapse recording. With the former two microscopes, 37°C/5% CO_2_ humid chambers were used for long-term observation.

### 2.17 Statistical Analysis

Biochemical and morphological data were quantified as described above. They were subjected to one-sided Welch’s *t-*test, one-way ANOVA, two-way ANOVA, and chi-squared test using Excel for Microsoft 365 (Microsoft) and Prism (v.7 and v.9.3.1, Graph Pad) software. For determining the peripheral distribution score (**Fig. 5C and F**), the signal intensity in the area within 5 μm from the cell periphery was divided by the total intensity in the cytoplasm.

### 2.18 Study Approval

The usage of hMSCs was approved by the Graduate School of Medicine, The University of Tokyo (approval No. 2023383NI). The recombinant DNA technology experiments were also conducted under the approval of the Graduate School of Medicine, The University of Tokyo (approval No. G24M0427).

## 3. Results

### 3.1 The Rab Antagonist CID1067700 Induces SHH-rich Secretomes

We stimulated bone-marrow-derived hMSCs overnight with DMSO (carrier alone), SF1670 (SF; the PI3K agonist), and CID1067700 (CID; the Rab antagonist). The secretomes were collected from the conditioned medium and characterized via immunoblotting. Secretomes from all the three treatments contained the sEV marker, TSG101 (**Fig. 1A**). However, only SF- and CID-secretomes contained SHH protein, unlike the DMSO-secretomes. The PI3K-dependency of SHH-EV secretion was consistent with our previous analysis using a polydactyly mouse model (Wang et al., 2022). CID treatment could also produce significant amounts of SHH-EVs in a shorter period, with lower levels of the sEV marker (2 h; **Supplementary Fig. S1**).

**Figure 1.**
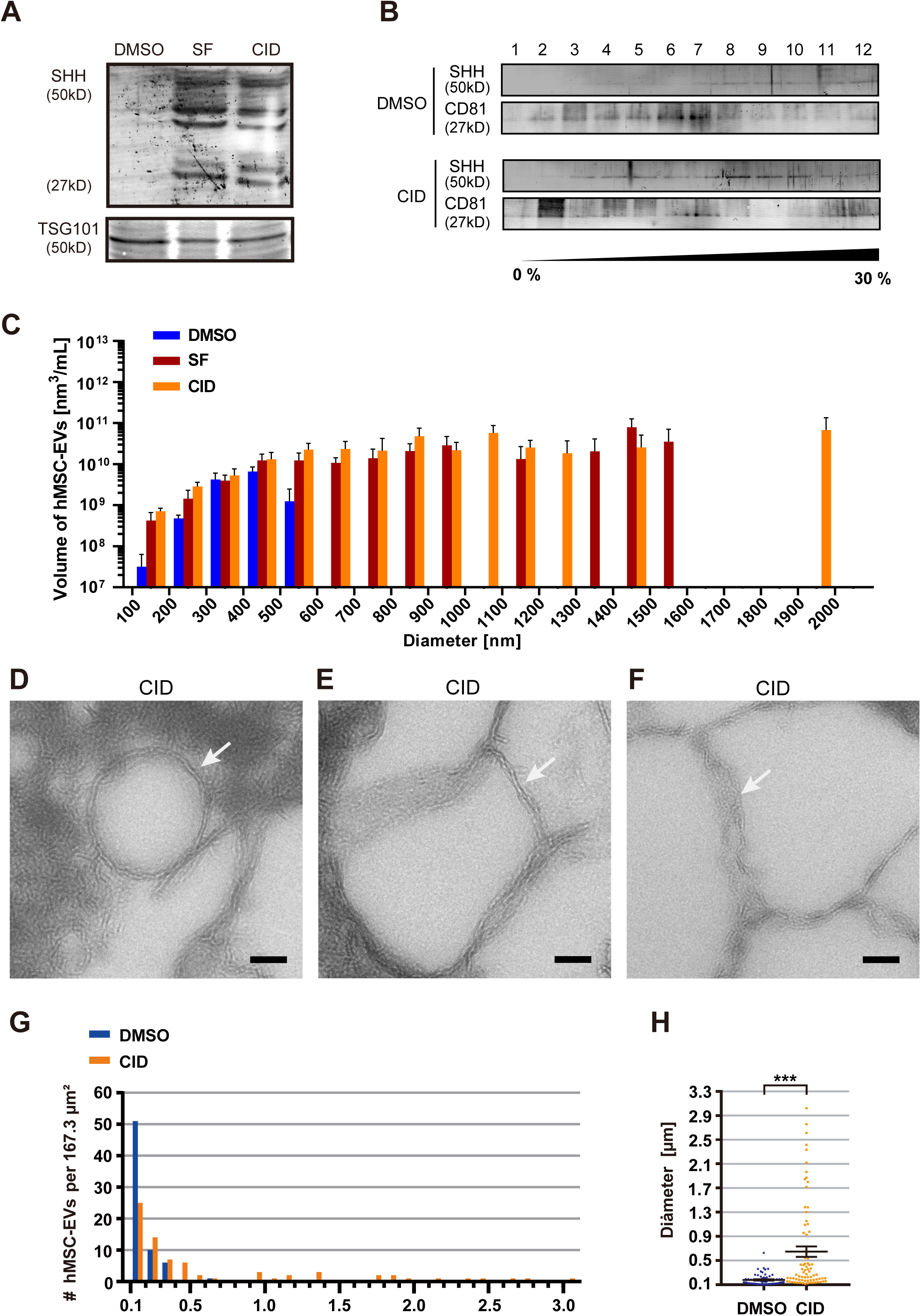
SF1670 (SF) and CID1067700 (CID) promote the secretion of sonic-hedgehog-containing large extracellular vesicles (SHH-lEVs) from human mesenchymal stem cells (hMSCs) (A) Immunoblotting of MSC-EVs, isolated after overnight treatment with 0.1% dimethyl sulfoxide (DMSO), 250 nM of SF, and 40 μM CID, using the indicated antibodies. Corresponds to **Supplementary Fig. S1**. (B) Fractions from the 0–30% Optiprep density gradient centrifugation of hMSC secretomes from DMSO- and CID-treated MSCs (40 μM, 24 h), probed against SHH and CD81. Note that SHH and CD81 were largely resolved in fractions 8–10 and 2–7, respectively. (C) Nanoparticle tracking analysis of hMSC-EVs, after overnight treatment with 0.1% DMSO, 250 nM of SF, and 40 μM CID, from three independent trials. Note that SF and CID facilitated the secretion of EVs with size >600 nm. Error bars, mean ± SEM. (D–H) Transmission electron microscopy of EV pellets from CID-treated hMSCs (40 μM, 24 h; D–F), accompanied by size distribution histograms (G and H). \*\*\**p* < 0.001; Welch’s *t*-test; *n* = 100.

To compare the buoyant density of SHH-lEVs and conventional sEVs, we performed density gradient centrifugation of the secretomes. CID treatment increased the production of SHH-lEVs, consistent with the immunoblotting results, and they were detected in heavier fractions than the conventional sEVs characterized by the CD81 marker (**Fig. 1B**).

Next, we analyzed the size profiles of the EVs via NTA of the conditioned medium (**Fig. 1C**). EVs larger than 600 nm were detected in SF- and CID-secretomes (SF, 3/3; CID, 3/3), but not in DMSO-secretomes (0/3; *p =* 0.01, chi-square test). However, EVs smaller than 600 nm were detected in all the secretomes (SF, 3/3; CID, 3/3; DMSO, 3/3). These results supported the notion that SF and CID facilitated the secretion of lEVs, especially those larger than 600 nm.

To morphologically characterize the CID-induced secretomes, we performed TEM of the EV pellets (**Fig. 1D–F**). Electron-lucent large particles, >600 nm in size, were significantly more abundant in CID-secretomes than in DMSO-secretomes. These large particles tended to be limited by lipid bilayers, which was indicative of their identity as EVs and not as lipoprotein particles or protein complexes. Size measurements revealed significant increase in lEVs after CID treatment (**Fig. 1G and H**), which was largely consistent with the NTA results. Collectively, these results indicated that CID treatment facilitates the biogenesis of lEVs. We obtained further evidence that these lEVs contained SHH via FACS analysis, as described in a later section.

### 3.2 CID-secretome Exhibits High Angiogenic Potency

We examined in vitro angiogenic potency of the secretomes using a bioassay employing HUVECs. Dissociated 6.5–8 × 10^4^ HUVECs were mixed with DMSO-, SF-, and CID-induced hMSC secretomes and incubated on Matrigel-coated chamber slides. These cells self-organize into vessel-like structures within 1–2 h and have been widely accepted as an in vitro experimental model of angiogenesis (Chance et al., 2020) (**Fig. 2A–C**). The levels of in vitro angiogenesis at 90 min after stimulation with the SF- and CID-secretomes were approximately 4–5 times of that achieved with the DMSO-secretome (**Fig. 2D**). These results indicated that the angiogenic potency of SF- and CID-secretomes is significantly higher than that of DMSO-secretome. As SHH protein is itself angiogenic (Renault et al., 2010), the observed tendency can primarily be explained by the enrichment of SHH in the lEVs in SF- and CID-secretomes.

**Figure 2.**
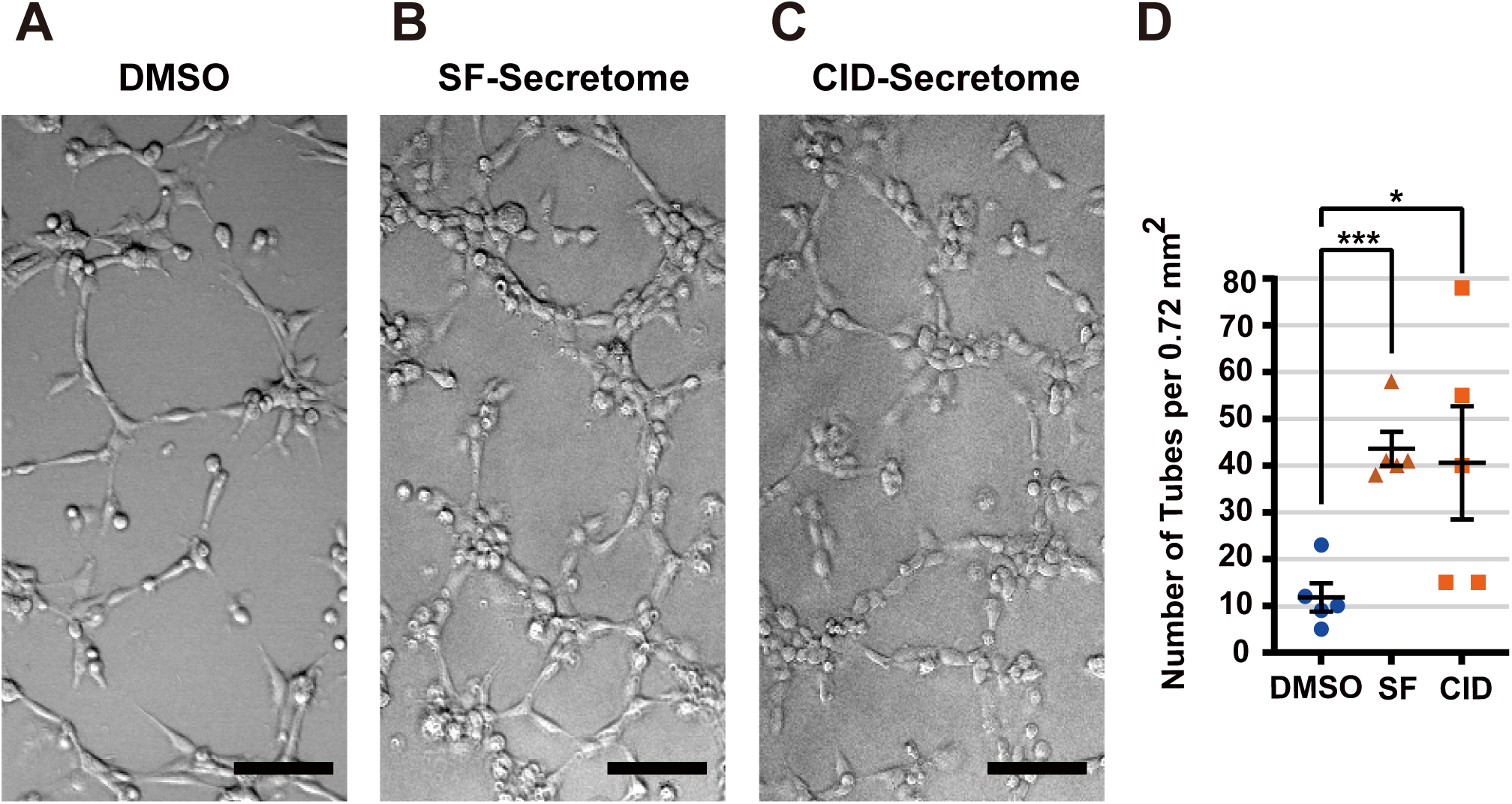
High angiogenic potency of SF1670 (SF)- and CID1067700 (CID)-secretomes. (A–C) Light microscopy of human umbilical vein endothelial cells (HUVECs) after 90 min of plating in 8-well chamber slides with the respective secretomes of human mesenchymal stem cells (hMSCs) in half of the conditioned medium in each 10 cm plate treated overnight with 0.1% dimethyl sulfoxide (DMSO), 250 nM of SF, or 40 μM CID. Scale bars, 100 μm. (D) Statistics of the density of tubes as an indicator of angiogenesis. \**p* < 0.05; \*\*\**p* < 0.001; one-way ANOVA; *n* = 5 independent experiments.

### 3.3 CID Facilitates SHH-lEV Secretion via Enrichment of Rab18-GDP

We transduced NIH3T3 fibroblasts with green fluorescent protein-tagged functional N-terminal fragment of SHH (SHH-N) (SHHN-EGFP; **Fig. 3A–E**). Green lEVs, approximately 1 μm in diameter (Wang et al., 2022), were observed to be moving in the culture medium with the counter flow, especially after CID treatment (**Fig. 3A**). We analyzed the density of the moving lEVs after different pharmacological treatments overnight.

**Figure 3.**
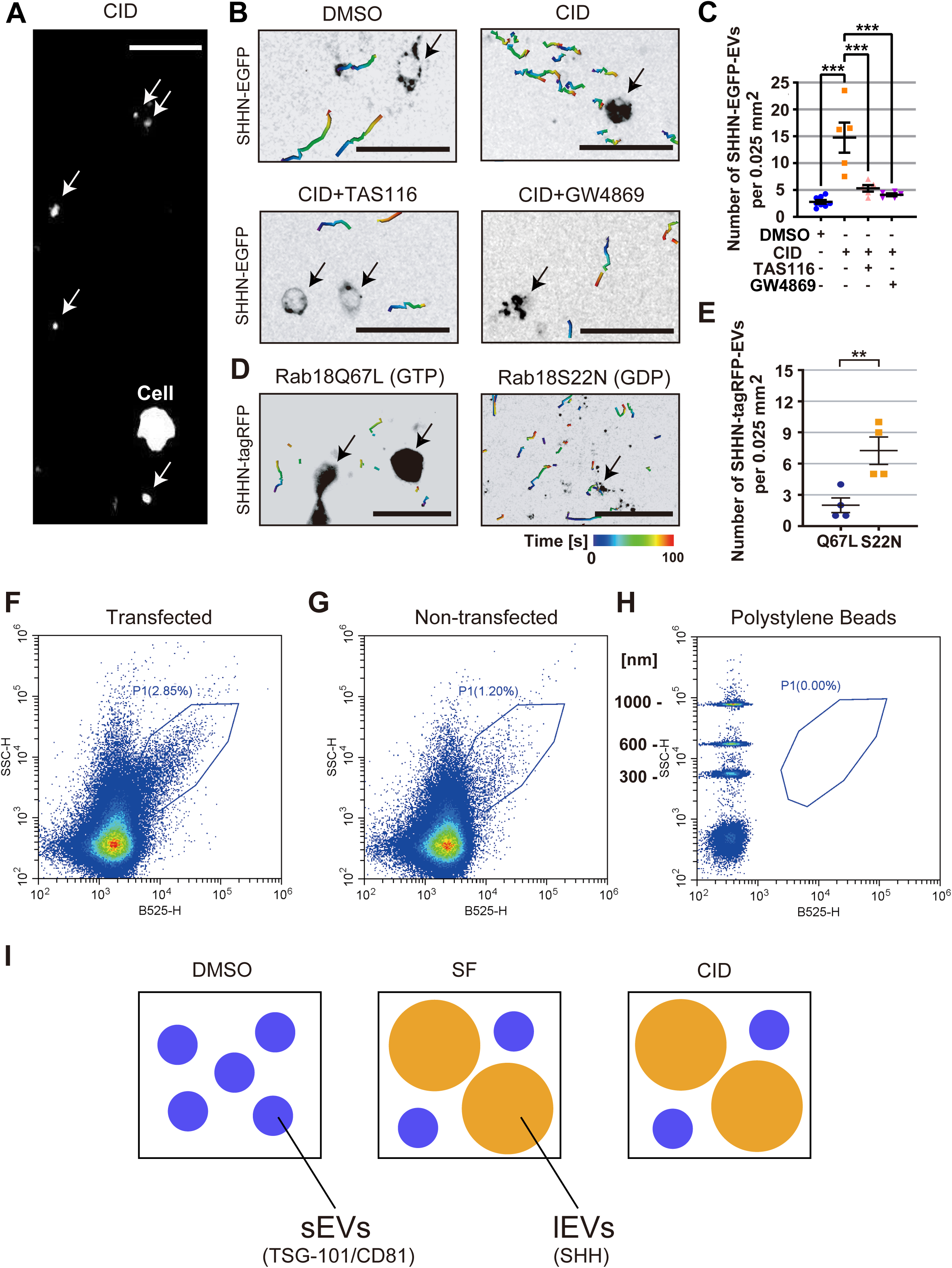
CID1067700 (CID) promotes the secretion of sonic-hedgehog-containing large extracellular vesicles (SHH-lEVs) (A–C) Fluorescence spinning disk microscopy of SHH-lEVs produced overnight by SHHN-EGFP-transduced NIH3T3 cells after treatment with 0.1% dimethyl sulfoxide (DMSO), CID (40 μM, 24 h), TAS-116 (0.5 μM, 48 h), and/or GW4869 (1.25 μM, 24 h); shown as a single frame (A) and as particle tracking images (B), around the transduced cells (Arrows), accompanied by the statistical analysis (C). Scale bars, 150 µm. Color bar, time sequence in s. The transduction efficiency of SHHN-EGFP was approximately 100%. \*\*\**p* < 0.001; one-way ANOVA; *n* = 5–8. Note that CID augmented the density of fluorescent particles, and that this effect was reversed by the Hsp90 inhibitor, TAS-116, or the nSMase2 inhibitor, GW4869. Corresponds to **Supplementary Movie S1**. (D and E) Fluorescence spinning disk microscopy images of SHHN-tagRFP-lEVs produced by the indicated Rab18-mutant-EGFP-stably expressing NIH3T3 cells doubly transduced with SHHN-tagRFP and incubated overnight after medium change, shown as particle tracking images (D) around the transduced cells (Arrows), accompanied by the statistical analysis (E). Scale bars, 200 µm. Color bar, time sequence in s. The transduction efficiency of SHHN-tagRFP was approximately 100%. \*\**p* < 0.01; Welch’s *t* test; *n* = 4. Note that Rab18-GDP augmented the density of fluorescent particles. Corresponds to **Supplementary Movie S2.** (F–H) FACS analysis of the conditioned medium of SHHN-mStayGold-expressing (F) and non-expressing (G) NIH3T3 cells that had been stimulated with 40 μM CID for 1 h (F and G) and that of polystyrene beads of the indicated sizes (H). *x* axis (B525-H), relative fluorescence intensity of the GFP channel; *y* axis (SSC-H), the relative size; P1, the gate for large fluorescent particles. Note that the SSC-H values for the beads were expected to be a little greater than those for EVs of the same size. The analysis was repeated twice. (I) Schematic representation of three kinds of mesenchymal stem cell secretomes containing small (sEVs; blue) and/or large extracellular vesicles (lEVs; orange).

CID treatment significantly augmented the density of SHH-lEVs in the culture medium (**Fig. 3B and C; Supplementary Movie S1**). This increase was reversed by simultaneous treatment with the Hsp90 inhibitor TAS-116 or with the nSMase2 inhibitor GW4869, which indicated that Hsp90 and nSMase2 activities are essential for CID-induced SHH-lEV biogenesis. This tendency was also reproduced by overexpressing nucleotide-state Rab18 mutants. We stably transfected fibroblasts with GTP- and GDP-bound mimetic Rab18-EGFP proteins. Notably, EGFP-Rab18Q67L(GTP)-transfected cells significantly retained the red signal of the SHHN-tagRFP protein in the cytoplasm, but the number of red lEVs in the medium was low. On the contrary, EGFP-Rab18S22N(GDP)-transfected cells showed a lower level of cytoplasmic red signal but significantly higher levels of red lEVs in the medium (**Fig. 3D and E; Supplementary Movie S2**). These data collectively suggested that Rab18-GDP significantly enhances the biogenesis of SHH-lEVs.

To directly measure the size of SHH-lEVs, we conducted FACS analysis of the conditioned medium of SHHN-mStayGold-transfected and non-transfected fibroblasts, which were simultaneously stimulated by CID for 1 h (**Fig. 3F–H**). The population of large and green fluorescent particles (Gate P1) increased approximately 2–3 folds upon transfection, providing evidence that the lEVs contained SHH (**Fig. 3I**). These data collectively suggested that CID enriches Rab18-GDP, which facilitates the biogenesis of SHH-lEVs in a manner dependent on Hsp90α and nSMase2.

### 3.4 SHH-lEVs are Secreted from the Perinuclear Region with Hsp90α Corona

We conducted time-lapse analyses of SHHN-EGFP-expressing fibroblasts for 1–2 h after CID treatment. Hsp90α-tagRFP was cotransfected to facilitate the dynamics. As depicted in spinning disk microscopy image with corresponding horizontal and lateral views (**Fig. 4A; Supplementary Movie S3**), SHHN-EGFP primarily accumulated in the perinuclear region by 30 min. The accumulation gradually approached cell surface from the perinuclear region via a vertical movement by 1 h.

**Figure 4.**
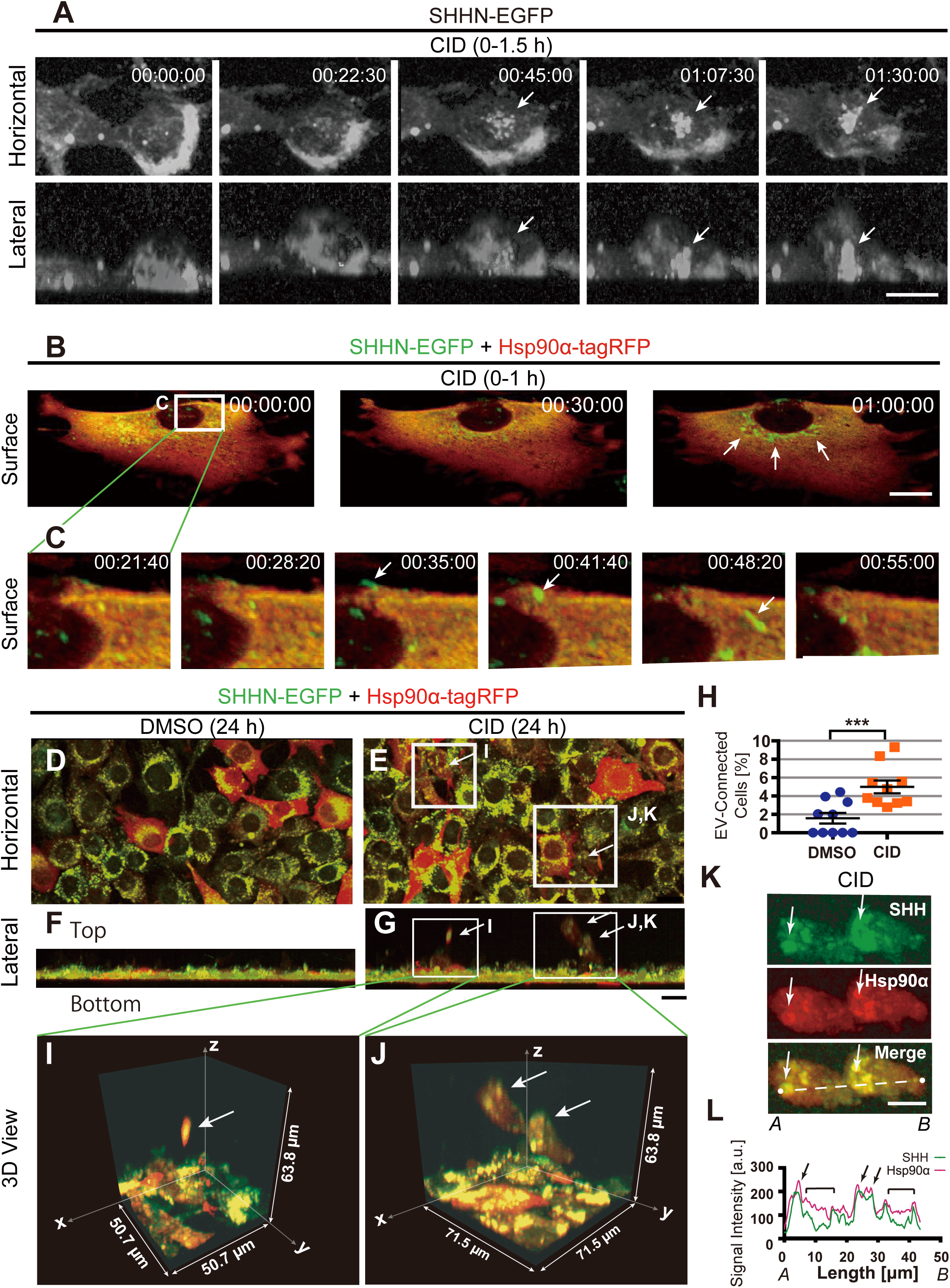
CID1067700 (CID) facilitates massive secretion of SHH and Hsp90α. (A) Three-dimensional (3D)-reconstituted time-lapse spinning disk microscopy images of SHHN-EGFP in NIH3T3 cells after double transfection with Hsp90α-tagRFP and stimulation with 40 μM CID for the indicated periods. The analysis was repeated three times. Scale bar, 20 µm. Arrows, SHH-lEV precursors vertically moving toward the cell surface. Time stamp, hh:mm:ss of the stimulation. Corresponds to **Supplementary Movie S3**. (B and C) Cell surface time-lapse images from four-dimensional (4D)-resonstruction of lattice lightsheet microscopy images of an SHHN-EGFP- and Hsp90-tagRFP-doubly transduced NIH3T3 cell under low-(B) and high-(C) magnification after the indicated periods of CID treatment (40 μM). The analysis was repeated twice. Scale bar, 20 µm. Arrows, fluorescent SHH-lEVs being secreted from the perinuclear region. Images acquired with a Lattice Lightsheet 7 microscope. Corresponds to **Supplementary Movie S4.** (D–L) Time-lapse spinning disk fluorescence microscopy images of NIH3T3 cells doubly transduced with SHHN-EGFP (green) and Hsp90α-tagRFP (red) and treated overnight with 0.1% dimethyl sulfoxide (DMSO) alone (D) or with 40 μM CID (D–J, K) in the corresponding horizontal (D and E), lateral (F and G), and 3D (I–K) views of the indicated areas in panel E, accompanied by statistical analysis of the fraction of cells with SHH-lEVs on cytonemes (H). (L) Intensity profiles in respective colors on the line *AB* in panel K. Reconstructions were performed using the Imaris 8 software. Scale bars, 20 μm. Arrows, SHH-lEVs being held in the Hsp90α corona associated with cytonemes. \*\*\**p* < 0.001; Welch’s *t*-test; *n* = 10. Corresponding to **Supplementary Movie S5**.

The secretion of SHH-lEVs from the cell surface was also observed using time-lapse lattice light sheet microscopy. Many tubulovesicular SHH-lEVs (1–2 μm) were secreted exclusively from the perinuclear region of SHHN-EGFP/Hsp90α-tagRFP doubly transfected fibroblasts by 1 h after CID treatment (**Fig. 4B**). We chased the secretory dynamics of an SHH-lEV in detail (**Fig. 4C; Supplementary Movie S4**). After secretion, it was associated with the cell surface for 15 min and suddenly slipped along the cell surface, disappearing from the optical field. This observation provided evidence that SHH-lEVs were secreted exclusively from the cell center.

We further observed the effect of long-term CID-treatment (**Fig. 4D–G**). After overnight incubation, we observed extracellular Hsp90α condensations, with diameters of 20–40 μm, containing SHH-lEVs that were floating in the medium, on cytoneme-like fibrous materials connected to the cell surface. The number of these extracellularly floating condensations was significantly increased by the CID treatment (**Fig. 4H**).

Based on high-magnification 3D views, these extracellular condensations could be broadly divided into two different classes of similar heights—one class had cilia-like straight protrusions (**Fig. 4I**) and the other was accompanied with cloud-like large Hsp90α condensations (**Fig. 4J; Supplementary Movie S5**). A higher magnification *z*-projection view of the latter type (**Fig. 4K**), with its linear profile of signal intensities (**Fig. 4L**), revealed that SHH-N/Hsp90α coaccumulating punctata (arrows, **Fig. 4K and L**) were surrounded by Hsp90α-dominant amorphous material (brackets, **Fig. 4L**). As these structures could be observed even after washing off the medium and fixing the cells, they may be cytoneme-bound EV corona consisting of Hsp90α polymeric condensates. We previously described a similar long fibrous interwoven structure associated with EVs in mouse embryonic ventral nodes (Tanaka et al., 2024), which may constitute a massive EV transfer system that systematically facilitates long-range movements between distinct cell populations.

### 3.5 SF1670 or CID Recruit Rab18 into Perinuclear Endosomes

We investigated the relevance of the Rab18 nucleotide cycle in SHH-lEV secretion. To test if PI3K signaling alters the distribution of Rab18, we treated Rab18-EGFP-stably transfected fibroblasts with the PI3K antagonist, LY294002, and the PI3K agonist or the PTEN antagonist, SF1670 (**Fig. 5A–C**). LY294002 facilitated the enlargement and scattering of Rab18 vesicles throughout the cytoplasm (**Fig. 5A**). However, SF1670 promoted their perinuclear clustering (**Fig. 5B**). Statistical analysis revealed that fluorescence intensity at the cell periphery after SF1670 treatment was significantly lower than that after LY294002 treatment (**Fig. 5C**), indicating that PI3K signaling significantly induces Rab18 clustering in the perinuclear region. To test if this occured via Rab18-GDP formation, we compared the distributions of the nucleotide-state-mimetic Rab18 protein (**Fig. 5D–F**). The accumulation of the Rab18S22N(GDP) protein was significantly more in the perinuclear region than that of the Rab18Q67L(GTP) protein. The apparent increase in puncta size might have been due to homophilic fusion between vesicles containing GTP-bound Rabs, as was reported for Rab5-GTP (Nielsen et al., 1999).

**Figure 5.**
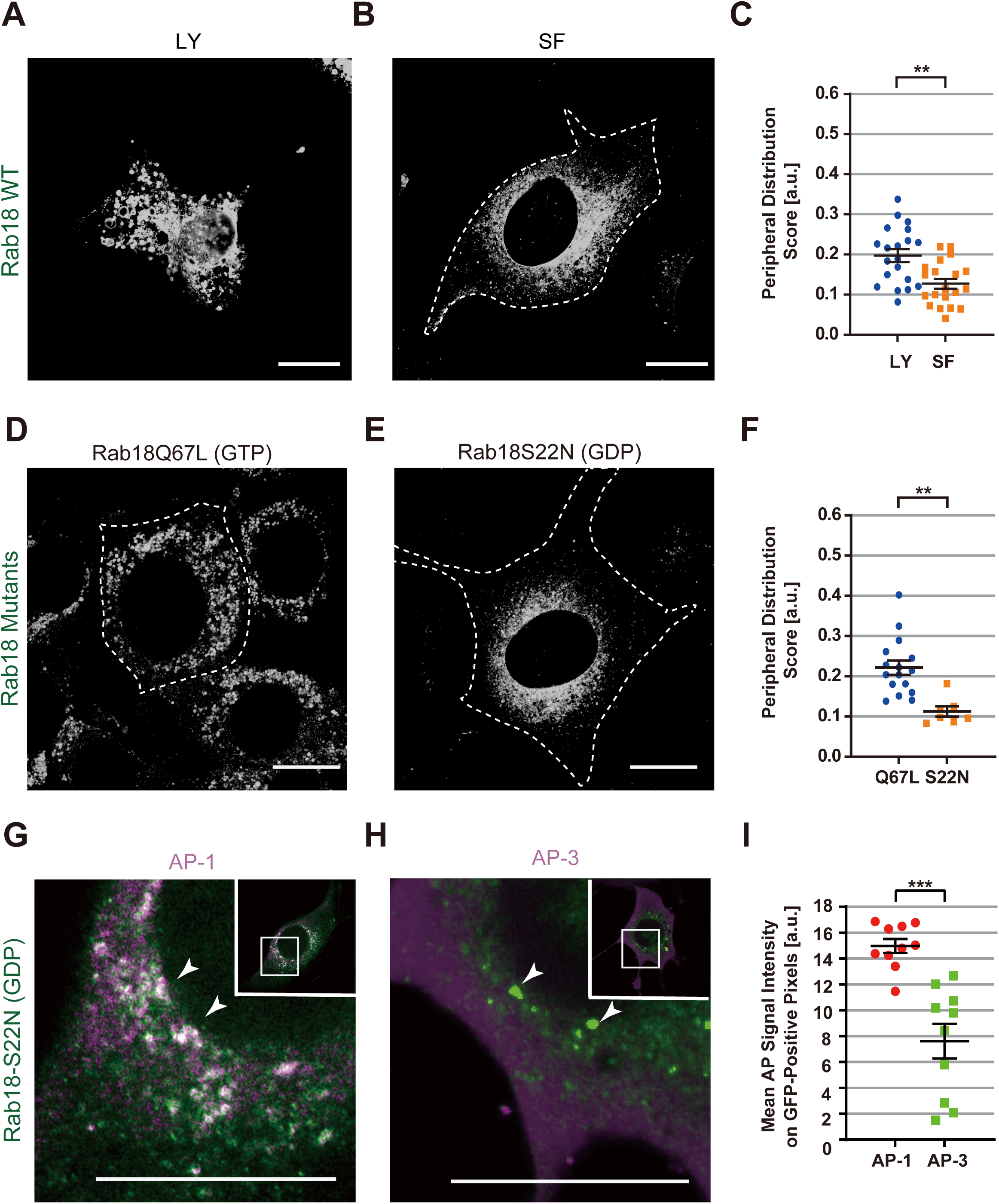
GDP-bound Rab18 facilitates the secretion of sonic-hedgehog-containing large extracellular vesicles (SHH-lEVs) (A–C) Fluorescence Airyscan microscopy of NIH3T3 cells stably transfected with SHHN-EGFP and treated with LY294002 (50 μM, 8 h; A) or SF1670 (250 nM, overnight; B), accompanied by the statistical analysis of the peripheral distribution scores (PDS), calculated as the ratio of the sum of fluorescence in an area, 5 μm from the cell periphery, and fluorescence over the whole cell (C). Scale bars, 20 μm. \*\**p* < 0.01; Welch’s *t* test; *n* = 20. (D–F) Fluorescence Airyscan microscopy of NIH3T3 cells stably transfected with EGFP-Rab18Q67L (D) and EGFP-Rab18S22N (E) that mimic the GTP- and GDP-bound states of Rab18, respectively; accompanied by the statistical analysis of the PDS (F). Scale bar, 20 μm. \*\**p <* 0.01; Welch’s *t-*test; *n* = 7–16. Scale bars, 20 µm. (G–I) Immunofluorescence Airyscan microscopy of NIH3T3 cells stably transfected with EGFP-Rab18S22N using antibodies against AP-1 (G) and AP-3 (H) with the colocalization statistics over the EGFP-positive pixels (I). Scale bars, 20 μm. Arrows, perinuclear Rab18S22N punctata. \*\*\**p* < 0.001; Welch’s *t* test; *n* = 10. Insets, the whole cell images. Scale bars, 20 µm.

To explore the identity of perinuclearly accumulated Rab18-GDP-containing organelles, we counterstained EGFP-Rab18S22N-transfected cells against the adaptor proteins AP-1 and AP-3 (**Fig. 5G–I**). AP-1 was more significantly colocalized with Rab18S22N-positive perinuclear vesicles than AP-3. Because AP-1 is involved in TGN-to-endosome sorting (Delevoye et al., 2009; Nakagawa et al., 2000; Schmidt et al., 2009), but AP-3 is involved in lysosomal biogenesis (Nakatsu and Ohno, 2003), the coaccumulation of Rab18-GDP and AP-1 in the perinuclear region suggested that Rab18-GDP accumulates in the endosomal system.

Overall, these results suggested that PI3K facilitates the enrichment of Rab18-GDP that accumulates into the perinuclear endosomes, consisting of a precursor for SHH-lEV biogenesis.

### 3.6 Rab18-GDP Endosomes Recruit Hsp90α and nSMase2

To further investigate the molecular mechanism for the facilitation of the maturation of SHH-lEVs by CID-induced Rab18-GDP enrichment, we conducted a glutathione-S-transferase (GST)-pulldown assay of mouse brain lysates against each nucleotide-state-mimetic Rab18 protein fused with GST (**Fig. 6A**). We found that Rab18-GDP was preferentially associated with Hsp90α and nSMase2. However, Rab18-GTP was associated with the kinesin-1 molecular motor KIF5, consistent with previous results suggesting that kinesin-1 cargo dynamics is partly regulated by Rab18 (Guadagno et al., 2020). Because kinesin and cytoplasmic dynein generate a balance in the tug-of-war for organelle distribution, the loss of a centrifugal motor upon enrichment of Rab18-GDP might facilitate the perinuclear clustering of Rab18-associated membrane organelles (Tanaka et al., 1998; Ueno et al., 2011). These differential binding preferences of Rab18 in each of the nucleotide states may be favorable for site-specific mechanism of the maturation of SHH-lEV precursors in the perinuclear region (**Fig. 6B**).

**Figure 6.**
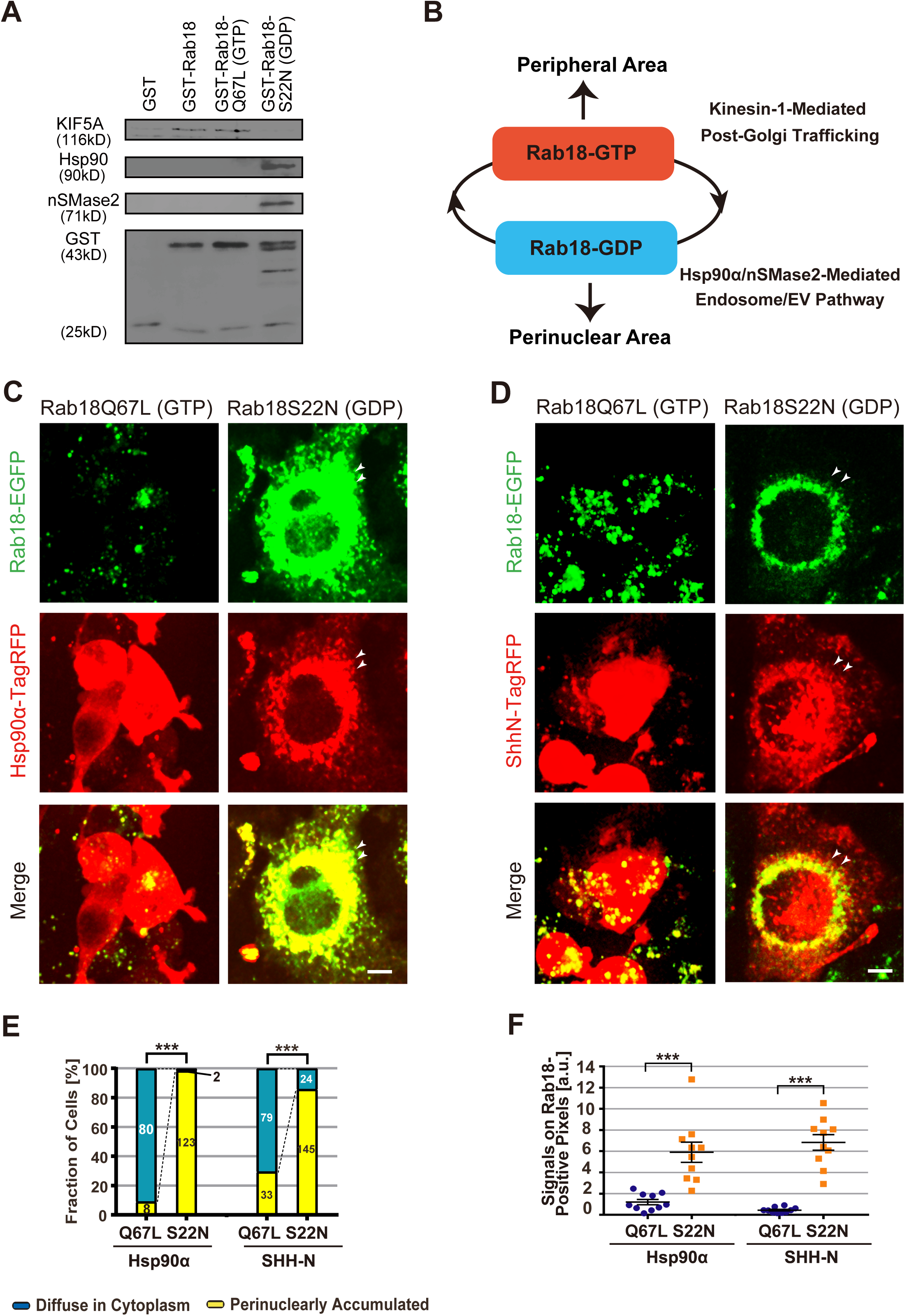
Hsp90α and nSMase2 are GDP-bound Rab18 effectors. (A) Immunoblotting following a pulldown assay of mouse brain lysates with the respective GST-tagged Rab18 mutant proteins, labeled with the indicated antibodies, and reproduced three times. Note that Rab18WT and Rab18Q67L (GTP-bound mimetic) were specifically associated with kinesin-1 (KIF5A), but Rab18S22N (GDP-bound mimetic) was associated with Hsp90α and nSMase2. (B) Schematic representation of the differential Rab18-GTP and -GDP effectors. (C and D) Double fluorescence spinning disk microscopy of NIH3T3 cells cotransfected with nucleotide-state mutants of EGFP-Rab18 (green; C and D) together with Hsp90α-tagRFP (red; C) and SHHN-tagRFP (red; D). Scale bars, 10 μm. Arrowheads, perinuclear coaccumulation of Rab18-S22N and its effectors. (E and F) Statistical analysis of perinuclear clustering of red signals (E) and colocalization of green and red signals (F) between Rab18-Q67L and -S22N-transfected cells corresponding to panels C and D. \*\*\**p* < 0.001; Chi-squared test; *n* = 88–125 cells (E); Welch’s *t*-test; *n* = 10 cells (F).

To test if the nucleotide cycle of Rab18 regulates the cargo distribution, we transduced tagRFP-labeled cargo molecules in the nucleotide-bound-state mimetic Rab18-EGFP cell lines (**Fig. 6C–F**). Hsp90α and SHH-N were highly coaccumulated with Rab18S22N (GDP-bound-mimetic) in perinuclear organelles. However, they were diffusely and abundantly distributed all over the cytoplasm of Rab18Q67L (GTP-bound mimetic) cells, without apparently showing punctate colocalization. Their cytoplasmic expression levels were significantly high, probably due to the impairment of cosecretion. These data collectively suggested that Rab18-GDP recruits Hsp90α and SHH into the perinuclear endosomes, which may be favorable for the extracellular secretion of lEVs through these organelles.

### 3.7 Rab18-GDP Enhances nSMase2-dependent Ceramide Biosynthesis

Because the Rab18-GDP effector, nSMase2, is a plasma-membrane-bound enzyme (Piacentino et al., 2022), we sought to determine if the association of Rab18-GDP augments its enzymatic activity in ceramide biosynthesis. We immunocytochemically analyzed the levels of ceramide in fibroblasts subjected to various pharmacological treatments.

CID treatment significantly augmented the ceramide levels, especially in the periphery of the cells (**Fig. 7A and B**). Notably, this increase could be reversed by cotreatment with GW4869, an inhibitor of nSMase2, which was further rescued by adding SF1670 (**Fig. 7C and D**). Because ceramide enlarges the endosomes and lysosomes (Li et al., 1999) and facilitates EV biogenesis (Fiorani et al., 2023; Horbay et al., 2022), the Rab18-GDP-mediated activation of nSMase2 may facilitate efficient secretion of lEVs. These data collectively indicated that PI3K–Rab18-GDP signaling induces an Hsp90- and nSMase2-mediated unconventional secretion of SHH-lEVs, which is exclusively from the perinuclear region (**Fig. 7E**).

**Figure 7.**
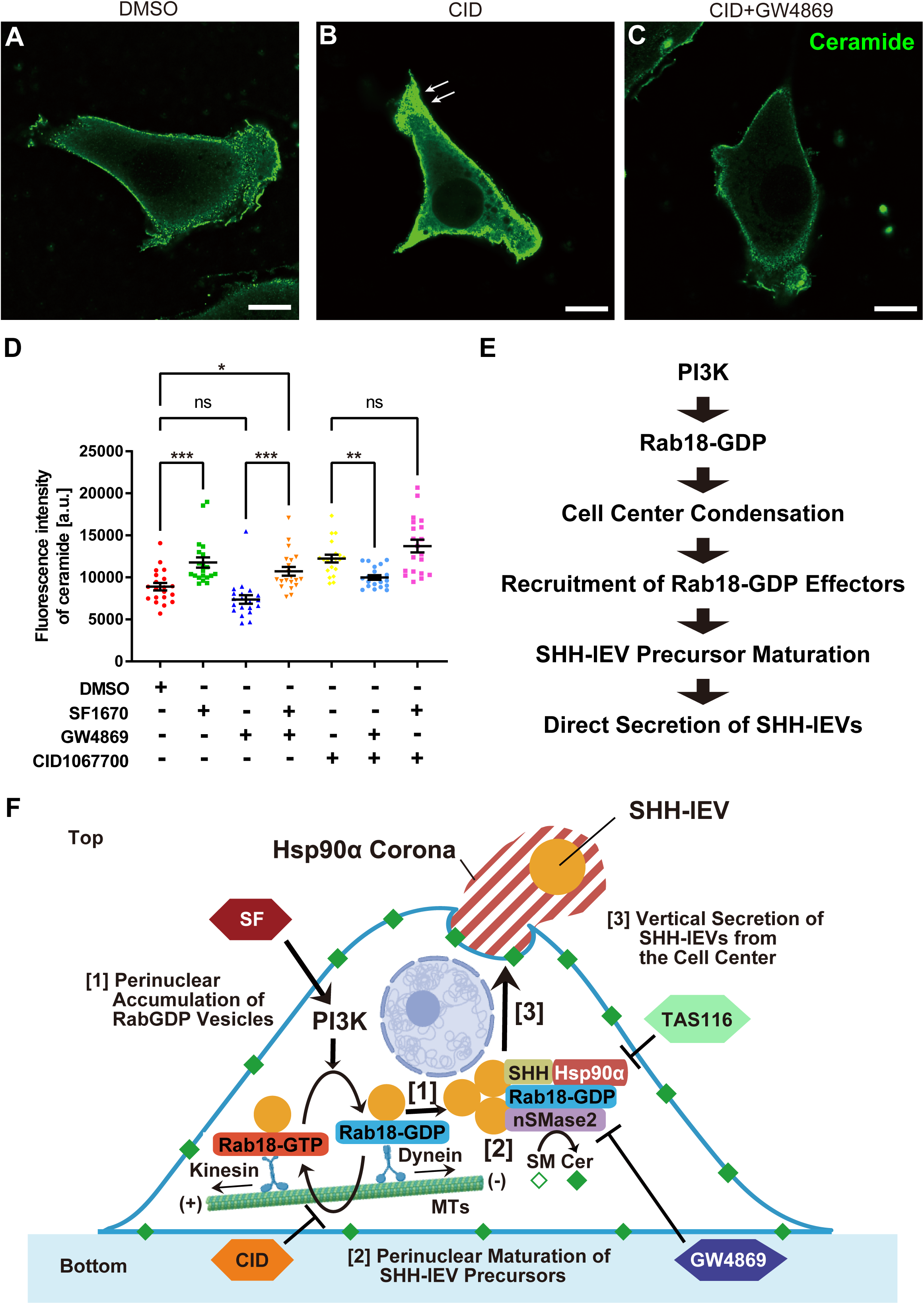
GDP-bound Rab18 promotes nSMase2-dependent ceramide biosynthesis. (A–D) Ceramide immunocytochemistry of NIH3T3 cells treated overnight with dimethyl sulfoxide (DMSO) alone (A), CID1067700 (CID) (40 µM; B); and CID plus the nSMase2 inhibitor GW4869 (1.25 µM, 24 h; C); accompanied by the statistical analysis (D). Scale bars, 10 μm. Images were taken using an LSM780-Airyscan microscope. Arrows, peripheral accumulation of ceramide. \**p <* 0.05; \*\**p <* 0.01; \*\*\**p* < 0.001; one-way ANOVA; *n* = 20. (E) Working hypothesis for the PI3K-mediated sonic-hedgehog-containing large extracellular vesicle (SHH-lEV) release pathway. (F) Schematic representation of the regulatory pathway and pharmacology of SHH-lEV secretion. Note the three steps of stimulation-secretion coupling specifically augmented by CID.

## 4. Discussion

In this study, we characterized angiogenic SHH-lEVs as large exosomes that are unconventionally secreted through the PI3K–Rab18-GDP signaling pathway (**Fig. 7F**). The strong angiogenic potency of these lEVs (**Fig. 2**) suggested their involvement in developmental and regenerative PI3K-mediated angiogenesis pathways (Kobialka et al., 2023; Ma et al., 2009). Because our data suggested that pharmacological treatments with either the PTEN inhibitor SF1670 or the Rab inhibitor CID1067700 significantly and specifically augmented the secretion of these angiogenic SHH-lEVs, they may have a clinical potential for utilization as an EV drug against a wide variety of vessel diseases, including thromboangiitis obliterans (TAO) and diabetic gangrene (Canha and Soares, 2023; Fujioka et al., 2023), and can be industrially produced. SHH-lEVs may also be used in regenerative medicine against intractable conditions, such as severe post-ischemic heart failure and spinal cord injury (Jia et al., 2021; Mackie et al., 2012). Accordingly, the present findings on the pharmacological pathways that stimulate the secretion of SHH-lEVs will facilitate translational research on lEVs.

Notably, the pan-Rab inhibitor CID1067700 (CID) that enriches the GDP-bound forms of Rabs specifically enhanced the secretion of SHH-lEVs and tended to suppress that of sEVs (**Fig. 1A; Supplementary Fig. S1**). This specific enhancement was mainly considered due to Rab18 inactivation because Rab18 is a major constituent of SHH-lEV precursors according to the data presented by us and other groups (Coulter et al., 2018; Wang et al., 2022), and overexpression of Rab18-GDP- and Rab18-GTP-mimetic proteins significantly enhances and suppresses the secretion of SHH-lEVs, respectively (**Fig. 5D–F**).

We propose that Rab18 is a smart master regulator of both the geometric and enzymatic activities of the SHH-lEV secretion machinery. Regarding the subcellular geometry, Rab18-GDP tended to accumulate in AP-1-positive endosomes in the perinuclear region, but Rab18-GTP tended to be dispersed throughout the cytoplasm (**Fig. 5D–F**). The centrifugal molecular motor, kinesin-1, exhibited a strong binding preference only for the GTP-bound form of Rab18 (**Fig. 6A and B**). This nucleotide-state-preference was consistent with that of other subtypes of kinesin motors (Niwa et al., 2008; Ueno et al., 2011). Accordingly, Rab18-GDP-vesicles may be dominantly associated with the centripetal motor, cytoplasmic dynein, which facilitates their perinuclear clustering due to a change in power balance in a tug-of-war situation.

PI3K stimulation enhanced the secretion of SHH-lEVs (**Fig. 1A**), consistent with a previous report (Wang et al., 2022). Because it significantly shifted the distribution of Rab18-EGFP toward the perinuclear region,PI3K signaling might provide the upstream signal to generate the GDP-bound Rab18 protein. This PI3K–Rab18 relationship is also supported by a previous report that insulin-mediated PI3K signaling similarly recruits Rab18 to the surface of lipid droplets in adipocytes (Pulido et al., 2011). As the canonical hedgehog signaling in the muscle cells facilitates the PI3K signal transduction (Elia et al., 2007), this PI3K–Rab18-GDP-mediated SHH-lEV secretion is also likely involved in a self-enhancing loop.

Regarding the regulation of enzymatic activity, Rab18-GDP was predominantly associated with Hsp90α and nSMase2 (**Fig. 6A**), both of which were essential for CID-stimulated SHH-lEV secretion based on pharmacological evidence (**Fig. 3B and C**). Hsp90α significantly enhanced the secretion of SHH-lEVs and was observed to be cosecreted with the SHH-lEVs (**Fig. 4**). This unconventional secretion paradigm of Hsp90α will provide additional insights into the Hsp90α-dependent mechanisms of the surface presentation of autoantigens (Srivastava, 2002). Because it facilitates the development of autoimmune diseases, it should be a future theme for research on unconventional secretion.

nSMase2 is an enzyme that converts sphingomyelin into the bioactive lipid, ceramide (Quadri and Bieberich, 2025), which augments EV biogenesis (Fiorani et al., 2023; Horbay et al., 2022). As the cellular levels of ceramide were significantly elevated by CID treatment (**Fig. 7**), the increase in the association of Rab18 with nSMase2 augmented its enzymatic activity. Accordingly, CID is suggested to enhance the Rab18-GDP-mediated enzymatic function that may facilitate the secretion of SHH-lEVs.

The perinuclear secretion of SHH-lEVs is distinct from that of sEVs containing CD81 or CD9, whose punctata are dispersed throughout the cytoplasm (Mathieu et al., 2021). Our data suggested that the Rab inhibitor CID only enhanced the secretion of SHH-lEVs (**Fig. 1A**). sEV secretion depends on both the GDP-bound form of Rab7 (Jiang et al., 2025) and the GTP-bound forms of Arl13b/Rab27a small molecular GTPases (Ostrowski et al., 2010; Verweij et al., 2022). Thus, sEV secretion likely consists of a dual process of (1) Rab-GDP-induced perinuclear clustering and (2) Rab-GTP/kinesin-induced centrifugal transport along the microtubules. This latter process in SHH-lEV secretion may be substituted by direct vertical elongation of the tubulovesicular SHH-lEV precursors, which may be driven by the contraction of actomyosin in the cell center (**Fig. 4A, Supplementary Movie S3**). Because of their huge size, they can reach the plasma membrane by slow massive extension, while their transport via the kinesin motors might be difficult. In previous research from our and other groups, loading SHH-N polypeptide to the cells and embryos could increase the size and/or efficacy of the secreted EVs (Coulter et al., 2018; Mackie et al., 2012; Tanaka et al., 2005; Wang et al., 2022). This may be partly because of the fact that lipid and cholesterol modifications of SHH-N polypeptide enhance its affinity for lipid bilayers (Lewis et al., 2001; Pepinsky et al., 1998). The high affinities between the SHH molecules (Whalen et al., 2013) may bridge the SHH-N-associated intraluminal vesicles of the multivesicular body. Accordingly, the SHH-N protein may facilitate homophilic fusion of its associated intraluminal vesicles in multivesicular bodies.

In conclusion, based on the findings of this study, we propose a PI3K–Rab18-GDP-mediated regulation that selectively enhances the unconventional secretion of SHH-lEVs. This mechanism of enhancement of SHH-lEV secretion via CID treatment that enriches GDP-bound Rab18, but not GTP-bound Rab18, was unexpectable from the previous notions on EV secretion (Blanc and Vidal, 2018). This efficient and selective strategy for the biogenesis of SHH-lEVs will be key to developing next-generation lEV therapeutics and should also enrich our understanding of the unconventional secretion of large particles.

## Supporting information

graphical abstract

## AUTHOR CONTRIBUTIONS

Y.T. directed and supervised the project and acquired the funding; S.W., R.I., and Y.T. conducted the experiments and wrote the paper.

## ACKNOWLEDGMENTS

We thank Marci Scidmore (NIH) for *EGFP-Rab18* expression vectors; Atsushi Miyawaki (RIKEN) for *mStaygold* expression vectors; Toshie Furuya (Univ Tokyo) for TEM observations; Yasushi Sakaguchi (DKSH Market Expansion Service Japan) for ZetaView analysis; Kanako Teshima (Beckman Coulter Japan) for FACS analysis; Fumiyoshi Ishidate (Kyoto Univ) for lattice lightsheet microscopy; and Shin-ichi Hasegawa (TWOCELLS, Co., Ltd.); Yiming Li, Shuzo Hasegawa, Yuya Kaneko, Maria Mulet Fernandez, Kyosuke Nakajima, Shimpei Goto, Momo Morikawa, Nobuhisa Onouchi, Takeshi Akamatsu, Hiromi Sato, Haruyo Fukuda, Nobutaka Hirokawa, Noboru Mizushima (Univ Tokyo); Yusuke Yoshioka, Takahiro Ochiya (Tokyo Med Univ); Ikuhiko Nakase (Osaka Metropolitan Univ); and Masahito Nakazaki (Sapporo Med Univ) for their technical help and valuable discussions. This work was supported by the GAP-Fund (8^th^, 12^th^, 14^th^ terms) from The University of Tokyo, by an AMED Grant jp22ym0126805 through the Translational Research Center, The University of Tokyo Hospital, and by KAKENHI Grant-in-Aids 20K06634 and 25K09626 from JSPS, to Y.T.

## DATA AND MATERIALS AVAILABILITY

All raw data and materials are available upon request to the authors.

## SUPPLEMENTARY MOVIE LEGENDS

**Movie S1. CID1067700 (CID) induces the secretion of sonic-hedgehog-containing large extracellular vesicles (SHH-lEVs) into the culture medium**

Fluorescent time-lapse spinning disk microscopy of NIH3T3 cells transduced with a SHHN-EGFP expression vector under the indicated pharmacological conditions, showing facilitation of SHH-lEV secretion by CID (40 µM, 24 h), which was antagonized by the Hsp90 inhibitor TAS116 (0.5 μM, 48 h) and by the nSMase inhibitor GW4896 (1.25 μM, 24 h). The entire movie corresponds to 100 s. Scale bar, 50 μm. Corresponds to **Fig. 3B**.

**Movie S2. GDP-bound Rab18 induces the secretion of sonic-hedgehog-containing large extracellular vesicles (SHH-lEVs)**

Fluorescent time-lapse spinning disk microscopy of NIH3T3 cells transduced with a SHHN-tagRFP expression vector together with the indicated EGFP-tagged Rab18 nucleotide-state-mimicking mutants (S22N for GDP-bound and Q61L for GTP-bound forms). The red channel images are shown. Note that the SHH-lEV secretion was facilitated by Rab18-GDP expression, but cytoplasmic accumulation of SHH was enhanced by Rab18-GTP expression. The entire movie corresponds to 100 s. Scale bar, 50 μm. Corresponds to **Fig. 3D**.

**Movie S3. CID1067700 (CID)-induced SHH vesicle motility in two successive phases**

Fluorescent time-lapse spinning disk microscopy of NIH3T3 cells transduced with SHHN-EGFP/Hsp90-tagRFP expression vectors. The green channel images are shown. Note that the horizontal perinuclear accumulation of SHHN-EGFP-carrying membrane organelles (arrow) is followed by a vertical movement toward the plasma membrane. The whole movie corresponds to 1.5 h. Scale bar, 20 μm. Corresponds to **Fig. 4A**.

**Movie S4. CID1067700 (CID)-induced the secretion of sonic-hedgehog-containing large extracellular vesicles (SHH-lEVs) from the perinuclear region**

Fluorescent time-lapse lattice lightsheet microscopy of the surface of an NIH3T3 cell transduced with SHHN-EGFP (green)/Hsp90α-tagRFP (red) expression vectors. Both channel images are shown. Note a green lEV, measuring 1–2 μm, secreted from the perinuclear region. The entire movie corresponds to 1 h. Scale bar, 20 μm. Corresponds to **Fig. 4B**.

**Movie S5. Hsp90α corona carrying sonic-hedgehog-containing large extracellular vesicles (SHH-lEVs) is induced by long-term CID1067700 (CID) treatment**

Rotation view of three-dimensional-reconstructed *z-*stack images of NIH3T3 cells cotransduced with SHHN-EGFP (green) and Hsp90-tagRFP (red), associated with extracellular condensations above the cell layer. Scale bar, 30 μm. Corresponds to **Fig. 4J**.

