## Supplementary figures and images for "Unconventional secretion of angiogenic sonic-hedgehog-containing large extracellular vesicles is stimulated via the PI3K–Rab18-GDP pathway"

### graphical abstract

#### PI3K–Rab18-GDP-Induced SHH-lEV Secretion

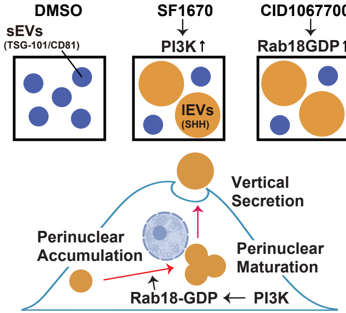
