## supplementary information for "Unconventional secretion of angiogenic sonic-hedgehog-containing large extracellular vesicles is stimulated via the PI3K–Rab18-GDP pathway"

Supplementary Fig. S1

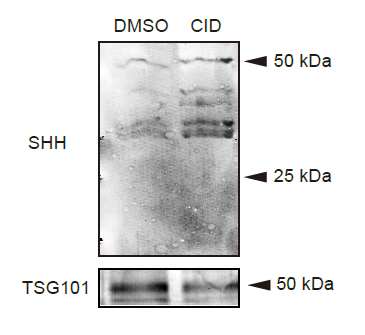

**Supplementary Fig. 1. CID induces SHH-lEV secretion by 2 h**

Immunoblotting of hMSC secretomes from culture supernatant after 2 h incubation with DMSO (carrier only) and 40 μM CID in EV-Up media. Note the increase in SHH but decrease in TSG101 levels by CID, largely consistent with the data after overnight incubation. Corresponds to **Fig. 1A**.
